# Drug repurposing based on a Quantum-Inspired method versus classical fingerprinting uncovers potential antivirals against SARS-CoV-2 including vitamin B12

**DOI:** 10.1101/2021.06.25.449609

**Authors:** Jose M. Jimenez-Guardeño, Ana Maria Ortega-Prieto, Borja Menendez Moreno, Thomas J.A. Maguire, Adam Richardson, Juan Ignacio Diaz-Hernandez, Javier Diez Perez, Mark Zuckerman, Albert Mercadal Playa, Carlos Cordero Deline, Michael H. Malim, Rocio T Martinez-Nunez

## Abstract

The COVID-19 pandemic has accelerated the need to identify new therapeutics at pace, including through drug repurposing. We employed a Quadratic Unbounded Binary Optimization (QUBO) model, to search for compounds similar to Remdesivir (RDV), the only antiviral against SARS-CoV-2 currently approved for human use, using a quantum-inspired device. We modelled RDV and compounds present in the DrugBank database as graphs, established the optimal parameters in our algorithm and resolved the Maximum Weighted Independent Set problem within the conflict graph generated. We also employed a traditional Tanimoto fingerprint model. The two methods yielded different lists of compounds, with some overlap. While GS-6620 was the top compound predicted by both models, the QUBO model predicted BMS-986094 as second best. The Tanimoto model predicted different forms of cobalamin, also known as vitamin B12. We then determined the half maximal inhibitory concentration (IC_50_) values in cell culture models of SARS-CoV-2 infection and assessed cytotoxicity. Lastly, we demonstrated efficacy against several variants including SARS-CoV-2 Strain England 2 (England 02/2020/407073), B.1.1.7 (Alpha), B.1.351 (Beta) and B.1.617.2 (Delta). Our data reveal that BMS-986094 and different forms of vitamin B12 are effective at inhibiting replication of all these variants of SARS-CoV-2. While BMS-986094 can cause secondary effects in humans as established by phase II trials, these findings suggest that vitamin B12 deserves consideration as a SARS-CoV-2 antiviral, particularly given its extended use and lack of toxicity in humans, and its availability and affordability. Our screening method can be employed in future searches for novel pharmacologic inhibitors, thus providing an approach for accelerating drug deployment.

## Introduction

The COVID-19 pandemic continues to cause high morbidity and mortality globally. Due to worldwide investment and international collaboration, multiple vaccines have been developed or are in the pipeline [1]. However, the ongoing emergence of new variants, different immunisation rates, supply chain issues, as well as the presence of smaller or larger outbreaks underlie the requirement for urgent treatments that can be rapidly deployed. Large outbreaks have overburdened hospitals worldwide due to the difficulty of both treating the disease and dealing with large numbers of patients. So far, therapies have focused on drugs that can improve the chances of survival during severe disease, with some antivirals [2] and antiviral candidates also emerging [3]. There is therefore an urgent need for pan-variant antivirals that are affordable, accessible and available worldwide.

From concept to treating a patient, it can take 10 years for a single treatment [1, 4]. Drug repositioning, repurposing, re-tasking, re-profiling or drug rescue is the process by which approved drugs are employed to treat a disease they were not initially intended/designed for. The main strategies are based on known pharmacological side-effects (e.g. Viagra [5]), library drug screening *in vitro* or computational approaches. The latter offers an advantage as processes can be modelled and investigated *in silico*, which allows for higher throughput than *wet lab* experiments. Virtual screening can be based on genetic information about the disease mechanisms, similarity with other diseases for which the drug is intended for, biological pathways that are common and/or known to be affected by certain drugs, or molecular modelling. Within the latter, molecular docking is perhaps the most common, where structures of targets are screened against libraries of compounds that will fit or *dock* into relevant sites [6].

Virtual screening has therefore become essential at the early stages of drug discovery. However, the process still typically takes a long time to execute since it generally relies on measuring chemical similarities among molecules, mainly to establish potential interactions between enzyme-substrate or receptor-ligand. Even for today’s processors, this exercise comprises a major challenge since it is computationally heavy and expensive. Accordingly, most of the well-known methods typically use 2D molecular fingerprints to include structural information that represents substructural characteristics of molecules as vectors. These methods do not take into consideration relevant aspects of molecular structures such as 3D folding, although they are efficient in terms of execution times. At the expense of higher computing times, considering 3D structural properties of molecules increases the accuracy of results [7]. 3D information from a given molecule can be encoded as a graph. In order to calculate the similarity between molecules, a new graph that contains information regarding the two molecules is required, allowing for better and faster comparisons to solve an optimization problem known as the Maximum Independent Set (MIS) that extracts the similar parts of those two graphs. By using Quantum or Quantum-inspired Computing, the mathematical model is able to manage this kind of information while having shorter execution times, up to 60 times faster.

The Randomised Evaluation of COVID-19 Therapy (RECOVERY) trial is an exemplar of drug repositioning during COVID-19: a multi-center trial that allows investigating the effectiveness of approved drugs for COVID-19 treatment. Tocilizumab and dexamethasone have been shown to improve survival in hospitalized patients [8–10] and are now used in the clinic. Remdesivir (RDV), initially designed against Ebola virus for which it failed to show efficacy in human trials [11] has also been approved for use in COVID-19 patients [2]. RDV is currently the only antiviral drug approved for use against SARS-CoV-2 infection albeit conflicting recent data [12]. Its action relies on its properties as a nucleoside analogue, whereby it binds to the RNA-dependent polymerase (RdRP) of SARS-CoV-2 and inhibits chain elongation [13]. Molnupiravir is another nucleoside analogue that is emerging as a potential antiviral against SARS-CoV-2 [3]. Although these compounds exhibit specific effects on viral, but not human, polymerases, there are multiple side effects associated with them, such as nausea or hepatic impairment [14]. Moreover, they are costly and thus implementation requires economic efforts that are unaffordable in many countries and settings. There is therefore an urgent need to identify novel antiviral compounds that exhibit low to no side effects, and that are readily and economically available.

We set out to investigate novel SARS-CoV-2 inhibitors based on the initial success of RDV. We modelled RDV as a graph and then screened the DrugBank dataset for compounds already approved for human use, employing a Quadratic Unbounded Binary Optimization (QUBO) model that runs on a quantum-inspired device, as well as a more traditional fingerprint method, the Tanimoto index [15] that runs on a regular laptop. Both algorithms predicted several candidates with high similarities to RDV, with GS-6620 being predicted by both. QUBO predicted BMS-986094 as second best and Tanimoto several forms of cobamamide, also known as vitamin B12. We then used cultured cell assays to determine the SARS-CoV-2 inhibitory capabilities of these compounds. BMS-986094, hydroxocobalamin, methylcobalamin and cobamamide all proved effective, with an inhibitory concentration to reduced infection by half (IC_50_) in ranges that are suitable for human use. Finally, we showed that these compounds are effective against a number of SARS-CoV-2 variants, including those known as Alpha, Beta and Delta (B.1.1.7, B.1.351 and B.1.617.2, respectively) in two cellular models. Our data illustrate the power of employing quantum-inspired computing for drug repurposing as well as a potential new role for vitamin B12 in treating SARS-CoV-2 infections.

## Results

Our workflow is represented in Figure 1a. We firstly established a computational model to search for structurally similar drugs to RDV. We then assessed their effect on viral replication *in vitro* to establish IC_50_ values and determine cytotoxicity. Finally, we measured the antiviral effects in two cell lines and with a panel of SARS-CoV-2 variants.

**Figure 1.**
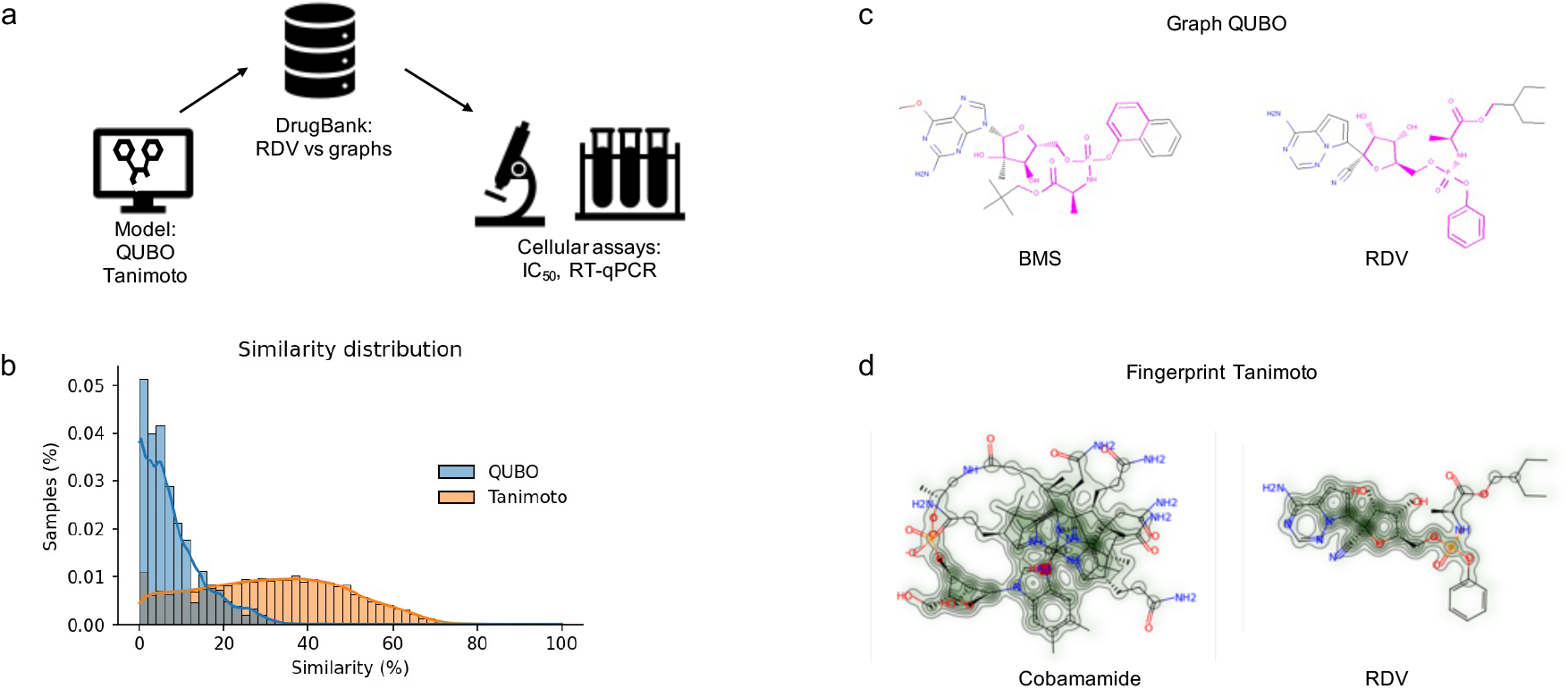
Molecular modelling. (a) pipeline employed in our project. RDV was firstly modelled as a graph and then screened against the DrugBank dataset. (b) Comparison of samples from the DrugBank predicted as similar to RDV by QUBO (blue) and Tanimoto (orange) models. (c) Graphic representation of BMS-986094 (BMS, left) and RDV (right) similarity according to QUBO. The magenta color represents the similar elements between the two molecules, including atoms as well as bonds, while the rest of the representation is the non-similar elements. d: Graphic representation of similarity for cobamamide (left) and RDV (right) generated by RDkit. The increasing green color represents more similarity between molecules.

### Molecular modelling

To search for similar compounds to RDV we firstly modelled chemical structures into graphs. A graph is a mathematical structure used to model pairwise relations between objects, where those objects are vertices in the graph and their relations are represented as edges. Similarity measures between two graphs can be derived from the Maximum Weighted Independent Set (MWIS) problem, with the MIS problem being a particular case in which all the weights are the same. The MIS problem is known to be a non-deterministic polynomial-time hard (NP-hard) problem [16]. A decision problem L is NP-hard if any problem in NP reduces to L [17]. Since these problems are difficult to approximate [18] more rapid computing approaches are required to solve NP-hard problems.

Considering structural properties of molecules to measure similarities among them increases the accuracy of results, but also increases higher computation times. With the commoditization of Quantum Computing in general and Quantum Annealers in particular, QUBO models have attracted attention as a way to describe a large variety of combinatorial optimization problems and thus can manage molecular structure information efficiently [16].

Fujitsu *Digital Annealer* [19] is a classical hardware inspired by Quantum Computing that solves these models in a fast, efficient way. We followed the three-step process described by Chams and colleagues [20] to solve the MWIS problem with *Digital Annealer*: firstly, acquire the model of the molecules as graphs, secondly generate a conflict graph to solve the MWIS problem, and thirdly measure the similarity between them considering the solution to the previous problem.

Atoms and ring structures were represented as vertices in a graph, while bonds connecting them were represented as edges that connect those vertices (atoms or rings). We considered the special case when two rings share one or more atoms; in this case, we created a new edge between those rings to formalize the graph structure. After creating the graph model for the two molecules being compared, we generated a new graph that gathered information about the molecules, or *conflict graph*.

In our conflict graphs, vertices represented possible matchings between the previous graphs, while edges represented conflicts between those new vertices. Matchings and conflicts directly depend on the given definition of similarity. As atoms can never be similar to rings and *vice versa*, the algorithm only compares atoms to atoms and rings to rings. For a comparison to be similar enough to enter as a vertex in the conflict graph, its value must be higher than a threshold value. In that case, a new vertex in the conflict graph is created containing the information of the two atoms or rings it refers to, as well as a measure of weight that depends on that information and the similarity value. We followed a similar process to construct the edges of the conflict graph. This rendered a new graph which ultimately had weights on the vertices and weights on the edges. Weights for the vertices indicated a positive value for the objective function of the optimization problem, while weights for edges indicated a negative value in the form of constraints (or penalizations to the model) for the optimization problem. We then constructed a QUBO model for solving the optimization problem given the conflict graph. This QUBO model was sent to *Digital Annealer* to solve the optimization problem and provide a solution. The stepwise algorithms are detailed in Materials and Methods.

The solution was a map that indicated the value 1 or 0 to every binary variable, meaning its presence or not in the independent set, respectively. Considering we also knew the map between each binary variable and the vertices from their respective molecules, we were able to calculate which atoms were similar and which were not. Detailed metrics are set out in Materials and Methods.

### Configuration of the algorithm: establishing *W*_*sim*_, *Min*_*sim*_, *W*_*edges*_

We employed a set of 100 instances for the preliminary experimentation to configure our algorithm. This set is described by Franco *et al* [21] and includes 100 pairs of molecules annotated by 143 experts that contain the SMILES (simplified molecular-input line-entry system, a line annotation that encodes molecular structures) for both molecules, the percentage of experts who determined they are similar and the percentage of experts who determined they are not similar. We assumed that if a percentage of experts annotated similarity between a pair of molecules, then that pair of molecules have a similarity value of that percentage of experts. We then tested the influence of different parameters that configure weights and thresholds to build the conflict graph within the algorithm (W_sim_, Min_sim_, W_edges_). We also tested the *δ* parameter that configures the similarity value.

W_sim_ is a similarity measure between two vertices that is tested against Min_sim_. Since other variables involved in this calculation, such as *vertices_similarity* and *edges_similarity*, are in the range [0, 1], we need to maintain the similarity measure of W_sim_ in that same range. However, we do not want to add the extreme values 0 and 1, as they represent similarity only among vertices (value 1) or edges (value 0). We therefore tested values for W_sim_ within [0.1, 0.9] in steps of 0.1. Min_sim_ is a threshold value for the similarity measure. Since that measure is in the range of [0, 1], Min_sim_ needs to be in that range too. We excluded pairs of vertices that did not have at least 50% similarity. Therefore, the final range of Min_sim_ was [0.5, 1], tested in steps of 0.05. W_edges_ gives a weight that is added to the final weight depending on the similarity of adjacent vertices to the vertices being compared. The value for W_edges_ was in the range of [0, 12], which was tested in steps of 1. The higher the value, the more importance the algorithm is giving to the similarity among adjacent vertices. The *δ* parameter also ranges from 0 to 1, and we test values in steps of 0.1. We tested these values and recorded the minimum and maximum similarity values prior to computing the final similarity value depending on the *δ* parameter.

W_sim_, Min_sim_ and W_edges_ influence the behavior of the algorithm and thus affect the result of the QUBO model for which we wanted to combine all three. This means we had 9 different values for W_sim_, 11 different values for Min_sim_, and 13 different values for W_edges_, totalling 1287 different combinations. The quality of the solutions was calculated as the Maximum Error (ME) of the similarity measure given by the QUBO model compared against the similarity measure given by the experts for each pair of molecules, averaged over all the results for each value of each parameter. We used the ME as a metric since it captures the worst-case error between the value given by the model and the value given by the experts. We thus considered the value that minimized the ME for each parameter for the final model. For each parameter we report a summary of the quality of the solutions considering the value of the parameters (Tables 1–4 for W_sim_, Min_sim_, W_edges_ and *δ*, respectively), where we have highlighted in bold the best values.

**Table 1.** Results of the preliminary experiment for W_sim_ parameter.

| $W_{sim}$ value | Max Error |
| --- | --- |
| 0.1 | 37.97 |
| 0.2 | 37.5 |
| **0.3** | **37.3** |
| 0.4 | 38.06 |
| 0.5 | 39.94 |
| 0.6 | 42.56 |
| 0.7 | 42.7 |
| 0.8 | 42.7 |
| 0.9 | 42.27 |

**Table 2.**
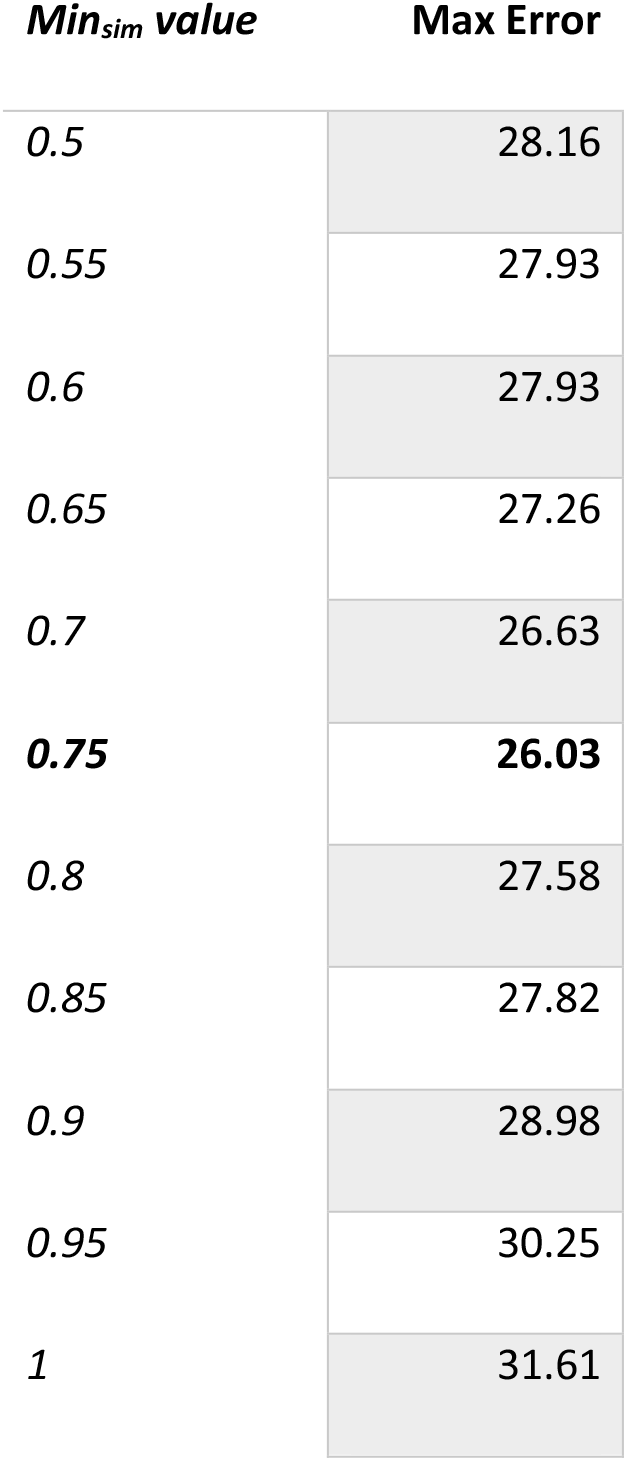
Results of the preliminary experiment for Min_sim_ parameter.

| $Min_{sim}$ value | Max Error |
| --- | --- |
| 0.5 | 28.16 |
| 0.55 | 27.93 |
| 0.6 | 27.93 |
| 0.65 | 27.26 |
| 0.7 | 26.63 |
| **0.75** | **26.03** |
| 0.8 | 27.58 |
| 0.85 | 27.82 |
| 0.9 | 28.98 |
| 0.95 | 30.25 |
| 1 | 31.61 |

**Table 3.**
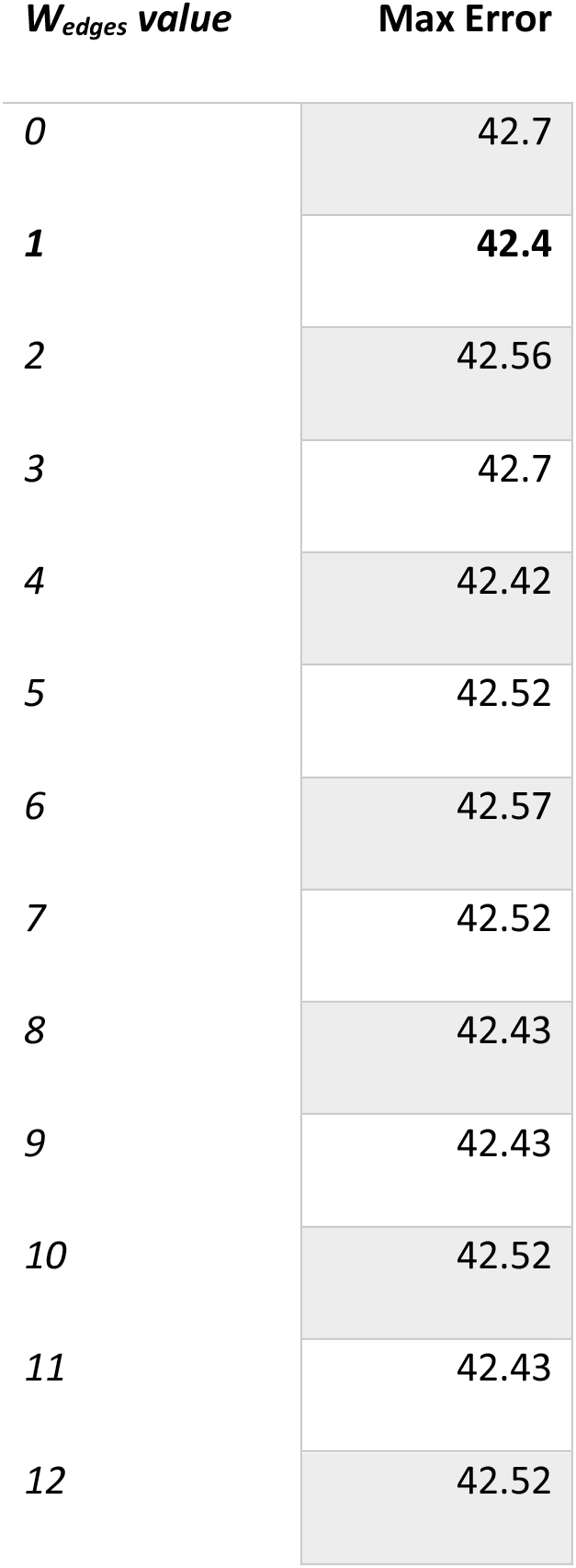
Results of the preliminary experiment for W_edges_ parameter.

| $W_{edges}$ value | Max Error |
| --- | --- |
| 0 | 42.7 |
| **1** | **42.4** |
| 2 | 42.56 |
| 3 | 42.7 |
| 4 | 42.42 |
| 5 | 42.52 |
| 6 | 42.57 |
| 7 | 42.52 |
| 8 | 42.43 |
| 9 | 42.43 |
| 10 | 42.52 |
| 11 | 42.43 |
| 12 | 42.52 |

**Table 4.**
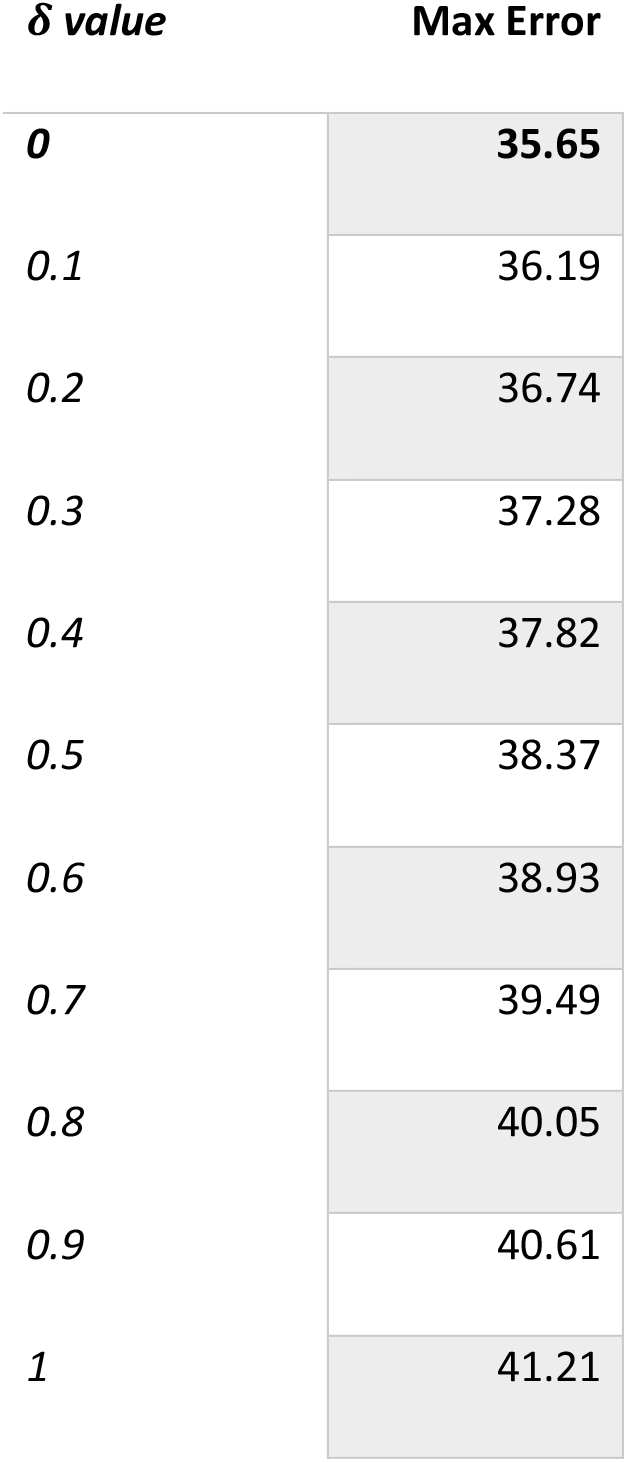
Results of the preliminary experiment for *δ* parameter.

| $\delta$ value | Max Error |
| --- | --- |
| **0** | **35.65** |
| 0.1 | 36.19 |
| 0.2 | 36.74 |
| 0.3 | 37.28 |
| 0.4 | 37.82 |
| 0.5 | 38.37 |
| 0.6 | 38.93 |
| 0.7 | 39.49 |
| 0.8 | 40.05 |
| 0.9 | 40.61 |
| 1 | 41.21 |

Table 4 shows the results for the *δ* parameter, for which the best ME is 0. The higher the value of *δ*, the higher the divergence, since the measure gives more weight to the maximum value of similarity. Thus, we selected 0.5 as the value for our algorithm, calculating then the similarity between two molecules as the average between the maximum and minimum values of similarity given by the model. Thus, the default configuration of the algorithm combined the values W_sim_ = 0.3, Min_sim_ = 0.75, W_edges_ = 1, and *δ* = 0.5.

### Comparison of molecules to Remdesivir

We applied the previously determined parameters of W_sim_, Min_sim_, W_edges_ and *δ* to our algorithm and searched for molecules with graphs similar to RDV. The set contained molecules approved for human use by the Federal and Drug Administration (FDA), following approval for Phase I or II clinical trials. In total, we compared 11405 compounds from the DrugBank to RDV. We also compared our outcomes with the ones given by a classical method based on fingerprints using RDKit [22, 23] and the Tanimoto measure [15]. We ran this method against the same dataset with the same target molecules. Figure 1b shows that the distribution of similarity values was different in the QUBO model compared to the Tanimoto method given by RDKit.

As seen in Tables 5 and 6, although GS-6620 came on top for both methods, most molecules predicted by both methods differed, illustrated by the different outputs of each method (Figures 1c and 1d). Figure 1c shows the representation of the second-best candidate for QUBO, BMS-986094 (BMS), where magenta represents similar elements between BMS, left, and RDV, right. The Tanimoto measure predicted as second best cobamamide, with the similarities (green) represented between cobamamide (left) and RDV (right) in Figure 1d.

**Table 5.** Top 10 molecules with similarity to Remdesivir according to QUBO.

| *Pairs* | **QUBO similarity** |
| --- | --- |
| *GS-6620* | 87.46 |
| *BMS-986094* | 61.49 |
| *Adafosbuvir* | 53.38 |
| *Sofosbuvir* | 51.59 |
| *Uprifosbuvir* | 51.59 |
| *Tenofovir alafenamide* | 48.7 |
| *Phenyl-uridine-5'-diphosphate* | 47.66 |
| *Thymectacin* | 47.14 |
| *Ethylhexyl methoxycrylene* | 46.63 |
| *Pantoyl Adenylate* | 44.05 |

**Table 6.** Top 10 molecules with similarity to Remdesivir according to the Tanimoto index.

| *Pairs* | **Tanimoto similarity** |
| --- | --- |
| *GS-6620* | 93.08 |
| *Cobamamide* | 80.03 |
| *Hydroxocobalamin* | 79.73 |
| *Mecobalamin* | 79.63 |
| *Heme D* | 79.27 |
| *Vintafolide* | 79.19 |
| *Thiostrepton* | 78.3 |
| *Vinflunine* | 78.22 |
| *Vincristine* | 78.16 |

### Assessment of IC_50_ and cytotoxicity of predicted compounds

We then evaluated the possible antiviral effects of the top predicted molecules by both methods. We did not assess GS-6620 as previous work has determined its lack of efficacy against SARS-CoV-2 [24]. We therefore assessed the antiviral effects of cobamamide (CB) and BMS and compared these with RDV in Vero E6 cells. Cells were incubated with serial dilutions of the compounds, infected with SARS-CoV-2 (England 02/2020/407073 isolate) and assessed for viral replication by plaque assay as well as cytotoxicity. Figure 2 demonstrates that all three compounds inhibited SARS-CoV-2 replication, with cobamamide showing an IC_50_ of 403μM, BMS of 26.6μM and RDV of 1μM. BMS appeared to be toxic to cells at >100μM, while cobamamide showed cytotoxicity of up to 20% at the highest dose of 1mM. RDV did not exert observable cytotoxic effects. Thus, our QUBO model is able to select for a highly efficient compound that inhibits SARS-CoV-2 replication *in vitro*.

**Figure 2.**
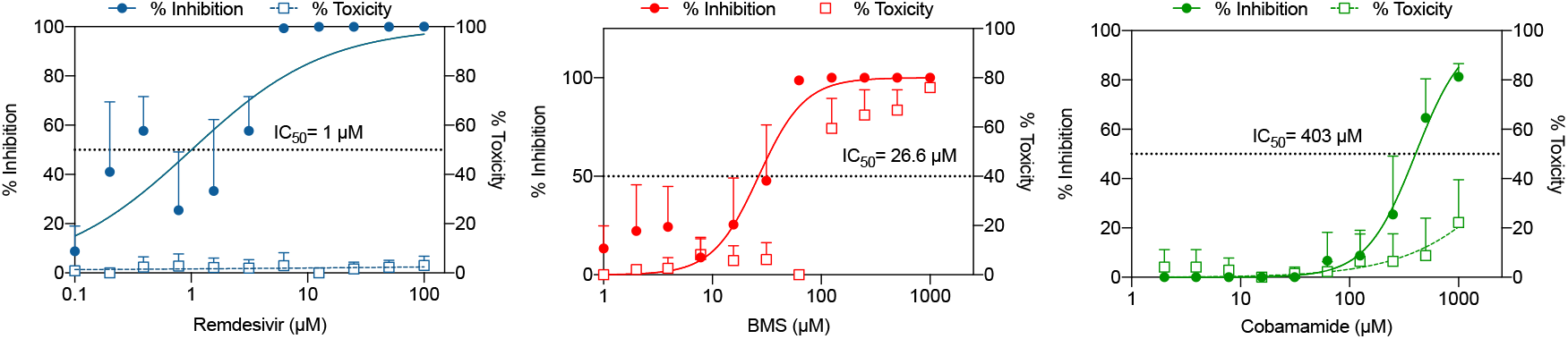
Dose-response inhibition of SARS-CoV-2 replication and cytotoxicity in Vero E6 cells. Vero E6 cells were treated with RDV, BMS or cobamamide at different concentrations for 2 h, followed by the addition of SARS-CoV-2 (MOI 0.05). After 1 h, cells were washed and cultured in compound-containing medium for 48 h. Virus production in the culture supernatants was quantified by plaque assay using Vero E6 cells. Cytotoxicity was measured in similarly treated but uninfected cultures via MTT assay. Data are mean ± s.d.; n = 3.

### BMS-986094 and several forms of vitamin B12 inhibit SARS-CoV-2 replication

We then assessed concentrations closest to their corresponding IC_50_ in both Vero E6 cells as well as Caco-2 cells, a human cell line permissive to SARS-CoV-2 infection. In addition to cobamamide, we also examined other forms of naturally occurring vitamin B12, namely methylcobalamin (MCB) and hydroxocobalamin (HCB). CB and MCB are two related corrinoid forms, which act as coenzymes in the mitochondria and cytosol, respectively, and differ in the R group of the central cobalamin. HCB is also abundant physiologically. They all share the core structure of cobalamine but differ in their upper ligands and are used as nutritional supplements [25]. Figure 3a shows that all compounds showed an antiviral effect, with a significant decrease in replication as measured by plaque assay both in Vero E6 cells (left) and Caco-2 cells (right).

**Figure 3.**
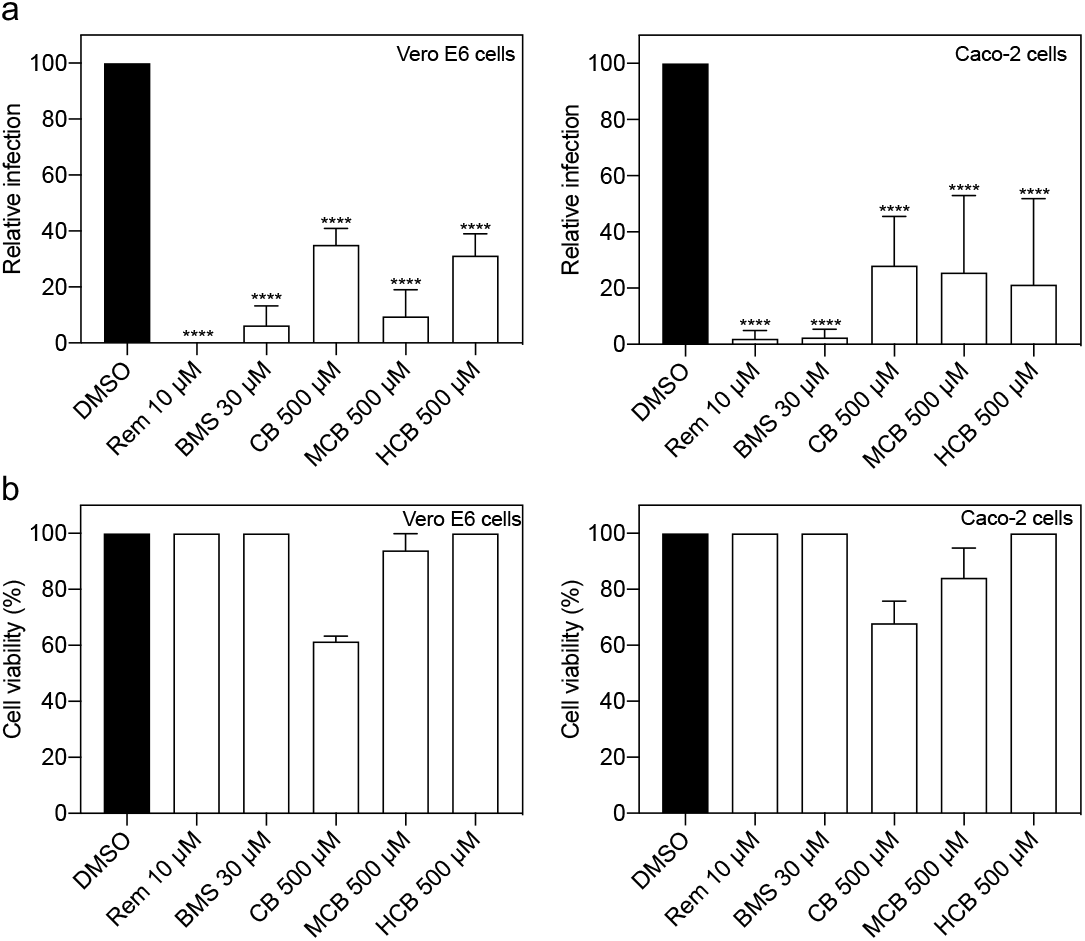
Inhibition of SARS-CoV-2 replication in different cell lines. (a) Vero E6 (n = 3) and Caco-2 cells (n = 4) were treated with DMSO, Remdesivir (Rem), BMS, cobamamide (CB), methylcobalamin (MCB) or hydroxocobalamin (HCB) at the indicated concentrations for 2 h, followed by the addition of SARS-CoV-2 (MOI 0.05 for Vero E6 and MOI 0.5 for Caco-2 cells). After 1 h, cells were washed and cultured in drug-containing medium for 48 h. Virus production in the culture supernatants was quantified by plaque assay using Vero E6 cells. (b) Cytotoxicity (n = 4) was measured in similarly treated but uninfected cultures via MTT assay. Data are mean ± s.d.; \*\*\*\**P* < 0.0001, ordinary one-way ANOVA with Dunnett’s multiple comparisons test.

These findings were recapitulated when we measured total and genomic viral RNA levels in cell lysates, with both consistently decreasing after exposure to BMS and different forms of vitamin B12 (Figure 4).

**Figure 4.**
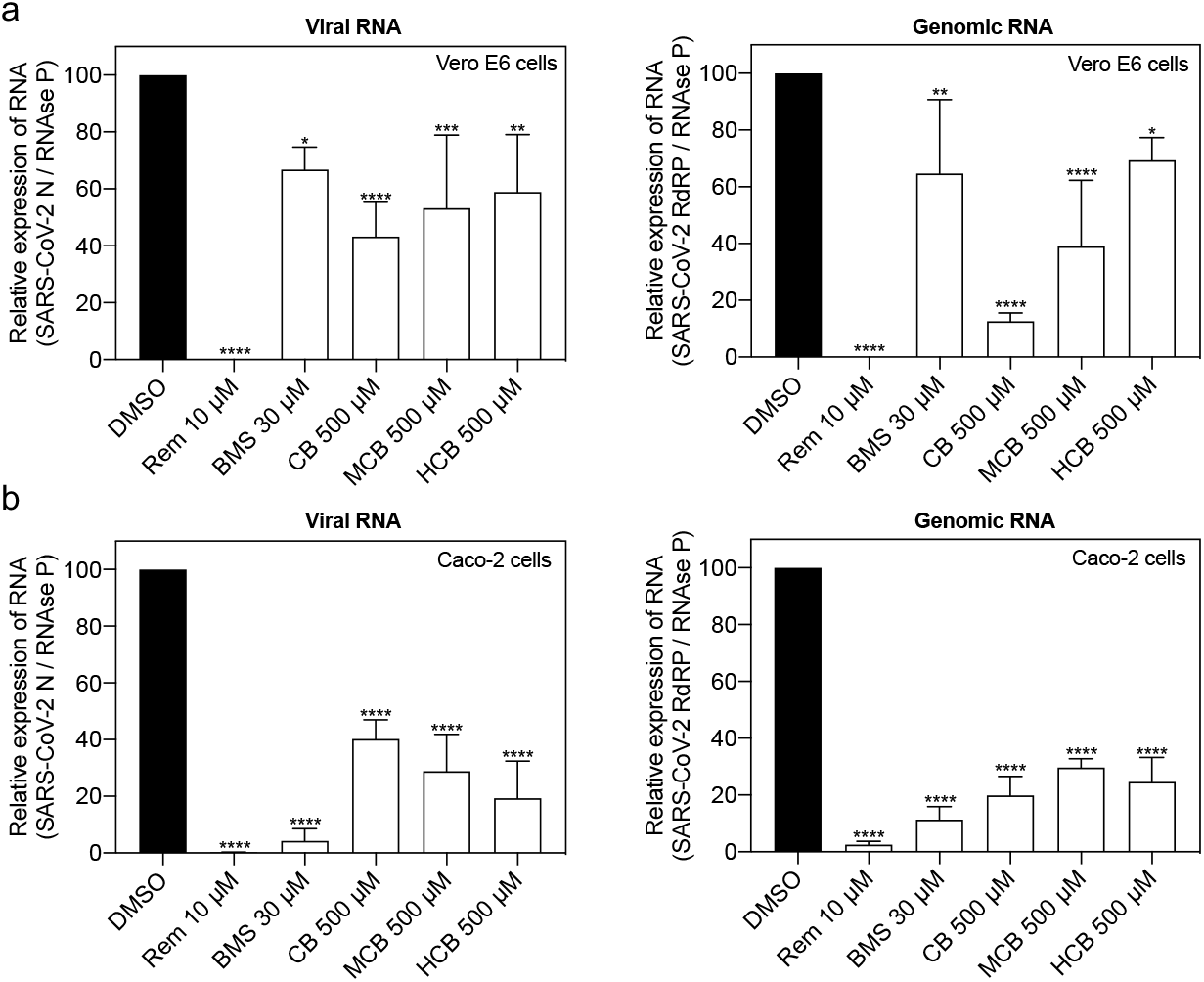
Identified compounds reduce SARS-CoV-2 RNA production in Vero E6 and Caco-2 cells. (a) Vero E6 (n = 4) and (b) Caco-2 cells (n = 3) were treated with DMSO, Remdesivir (Rem), BMS, cobamamide (CB), methylcobalamin (MCB) or hydroxocobalamin (HCB) at the indicated concentrations for 2 h, followed by the addition of SARS-CoV-2 (MOI 0.05 for Vero E6 and MOI 0.5 for Caco-2 cells). After 1 h cells were washed and cells lysates were collected 48 hours after infection. The levels of total and genomic viral RNA were analyzed with specific reverse transcription quantitative polymerase chain reactions. Data are mean ± s.d.; *P < 0.05, **P < 0.01, ***P < 0.001, ****P < 0.0001, ordinary one-way ANOVA with Dunnett’s multiple comparisons test.

### BMS-986094 and several forms of vitamin B12 inhibit viral replication of several SARS-CoV-2 variants

Given the continuing emergence of new variants of SARS-CoV-2, we also assessed the effects of BMS and vitamin B12 forms on the SARS-CoV-2 variants Alpha (B.1.1.7) and Beta (B.1.351) using two different isolates of each variant (Figure 5). We also assessed the effect on the recently identified Delta variant (B.617.2). Figures 5a and 5b show the effects of the different compounds on Vero E6 cells on Alpha and Beta variants, while Figures 5c and 5d show the effects on viral replication of these variants on Caco-2 cells. Assays for the new delta variant were also performed in Vero E6 and Caco-2 cells (Figure 5e, left and right, respectively). Consistent with earlier data, BMS and all forms of vitamin B12 suppressed the replication of this panel of SARS-CoV-2 isolates to similar degrees.

**Figure 5.**
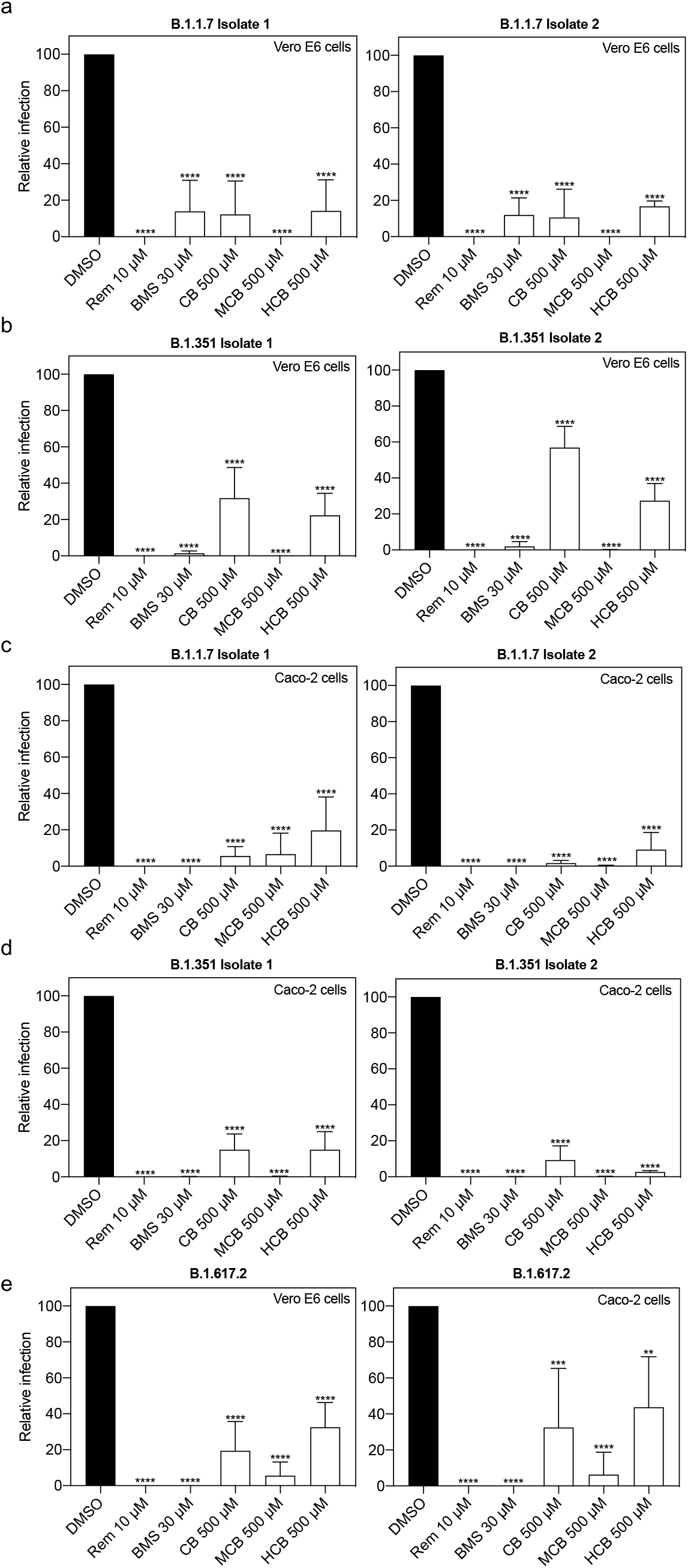
Effect of identified compounds at inhibiting SARS-CoV-2 replication is variant-independent. Vero E6 and Caco-2 cells were treated with DMSO, Remdesivir (Rem), BMS, cobamamide (CB), methylcobalamin (MCB) or hydroxocobalamin (HCB) at the indicated concentrations for 2 h, followed by the addition of SARS-CoV-2 variants isolates (a, c) B.1.1.7 (n = 3), (b, d) B.1.351 (n = 3) and (e) B.1.617.2 (n = 4) (MOI 0.05 for Vero E6 and MOI 0.5 for Caco-2 cells). After 1 h, cells were washed and cultured in drug-containing medium for 48 h. Virus production in the culture supernatants was quantified by plaque assay using Vero E6 cells. Data are mean ± s.d.; **P < 0.01, ***P < 0.001, ****P < 0.0001, ordinary one-way ANOVA with Dunnett’s multiple comparisons test.

## Discussion

We present a QUBO model employing the quantum-inspired device Fujitsu’s *Digital Annealer* [19] and a classical method based on fingerprints, Tanimoto measure [15], to seek compounds structurally similar to RDV in the DrugBank database (Figure 1a). We demonstrated how both approaches identified compounds that showed antiviral properties *in vitro* in two cell culture models (Figures 2–5). Our QUBO model rendered BMS-986094 as the second-best candidate (Table 5), which was proven to inhibit viral replication of several variants of SARS-CoV-2 (Figure 5). Importantly, we demonstrated that several forms of vitamin B12, namely cobamamide (adenosylcobalamin), methylcobalamin and hydroxocobalamin, currently used worldwide both orally and intravenously, inhibited replication of several variants of SARS-CoV-2 *in vitro* (Figure 5). Although animal models are required to determine whether these effects can be seen *in vivo*, there are already several studies investigating the possible relationship between vitamin B12 levels and SARS-CoV-2 infection outcome [26, 27].

Our approach to solve a MIS problem has been implemented in other settings. Chemical similarity approaches have their pitfalls: errors in chemical structures as well as physiological effects that exist beyond the structural relationship (for example, a metabolite of the original drug with a modified structure could be the active molecule) could limit the use of this approach in drug repurposing [28]. For example, Bollobás et al. [29] used MIS to help solve the Graph Coloring Problem. The authors extracted the maximal independent set of uncolored vertices iteratively to assign them the same color, repeating the process until the whole graph is colored. A similar technique was followed by Johnson [19], where several independent sets were first constructed and then a new color class selected after removing one of them with the smallest edge density. The algorithm repeated this process until a threshold number of uncolored vertices were left. After that, an exhaustive search algorithm was applied to color those remaining vertices. This problem was solved using several metaheuristics, such as Simulated Annealing [20], Tabu Search [30], GRASP [31], or Genetic and Hybrid Algorithms [32]. More recent studies included efficient ways based on fast local search procedures to produce near-optimal solutions to a wide variety of instances [33]. Jin *et al*. presented a general Swap-Based Local Search method based on Tabu Search to obtain the best-known solutions for two well-known set of instances used to evaluate algorithms [34].

Regarding the MWIS problem, one can implement a Genetic Algorithm [35], involving a two-fusion crossover operator and a mutation operator that is replaced by a heuristic-feasibility operator. The first operator considers both the structure and the fitness of two parent solutions to produce two children. The second one transforms infeasible solutions into feasible ones, with these being able to run in parallel. These problems involve determining DNA and amino acid sequence similarity and were reported by Joseph *et al* [36]. The authors proposed an algorithm that ran in O(n log n) computational time. With regards to QUBO, a more detailed description about how to formulate problems in this special mathematical way is described by Glover *et al* [37].

Recent studies showed the use of QUBO models to formulate the MIS problem to be solved by a Quantum Annealer. Similarity among a set of molecules has been implemented by relaxing the definition of measure by using the Maximum Co-k-plex relaxation method [38], a more general form of describing the MIS problem. Hernandez *et al* [39] reported that molecular similarity methods can take advantage of Quantum Annealers. The authors considered different relevant pharmacophore features to describe the molecules for ligand-based virtual screening, including atomic coordinates as features for comparison. The results showed better performance than fingerprint methods for most of the datasets used. Several attempts to solve combinatorial optimization problems with Fujitsu Digital Annealer were reported by Aramon *et al* [40]and Hong *et al* [41]. With regards to the choice of parameters for our QUBO model, we considered that the *δ* parameter value being 0 only described the nature of the instances and that similarity values had very different outcomes comparing the results from the experts from the ones given by the model.

Our data showed that the distribution of similarity values differed between the QUBO model vs the Tanimoto measure (Figure 1b, Tables 5 and 6), indicating that we could not directly compare solutions given by each method. Solutions that are good for one method may not be so in the other. Our experiments demonstrated that both approaches can discover potential antiviral compounds and that both models should be run in parallel (Figures 2–5). Solutions to the models (drug targets) produced in both cases can be very useful in different settings and show different properties in humans. BMS appeared to be a better candidate for SARS-CoV-2 replication inhibition (Figures 2–5) than vitamin 12 when we compared side-by-side IC_50_ concentrations. Previous studies have observed harmful cardiac effects, which halted its progression from phase II clinical trials [42]. The dose administered in the trial was 100mg orally (clinical trial NCT01629732). Our estimated IC_50_ of 26.6μM is higher than the established EC_50_ of 10nM for BMS in hepatitis C virus (HCV) infection [43], but further studies would be required to establish bioavailability in mucosae.

Our data demonstrated that several forms of vitamin B12 inhibit SARS-CoV-2 replication *in vitro* in two different cell lines (Figures 3–5). In humans, injections of up to 10mg have not shown side effects. The healthy range of vitamin B12 in blood is 118-701 pM. Vitamin B12 deficiency is treated with injections of 1mg hydroxocobalamin [44], with 1mg/mL hydroxocobalamin being 742.2μM. Higher doses of hydroxocobalamin are however tolerated. Upon cyanide poisoning, adults can receive up to 5g of hydroxocobalamin as an antidote intravenously, which would be well within the range of our calculated IC_50_ of 403μM. Moreover, previous studies on imaging show biodistribution of an intravenous In-111 labeled 5-deoxyadenosylcobalamin (AC) analog ([111In]AC) in nasal cavity and salivary glands [45]. There is also previous evidence of vitamin B12 having potential antiviral effects. An *in silico* screening by Narayanan and Nair [46] showed their second best docking score between methylcobalamin and nsp12 (the gene that encodes for RdRP). Vitamin B12 has been shown to improve outcome in hepatitis C virus (HCV) infection. HCV is a single-stranded RNA virus that has an internal ribosomal entry site (IRES) which interacts with cellular ribosomal subunits, and vitamin B12 has been reported to work by inhibiting HCV IRES-dependent translation [47]. Interestingly, BMS was originally designed against HCV.[48]

Our findings open the door to employing Quantum-inspired methods to inform drug repurposing. Our data showed novel compounds that were able to inhibit SARS-CoV-2 replication based on our QUBO model as well as the more traditional Tanimoto fingerprint, such as BMS and cobalamin derivatives. BMS warrants further investigation, while vitamin B12 is readily available from multiple sources, it is affordable, can be self-administered by patients, is available worldwide, and displays low-to-no toxicity at high doses.

## Materials and Methods

### Molecular modelling as graphs

Our *in silico* modelling was comprised of a three-step process: getting two molecules as graphs, solving a Maximum Independent Set problem with a Quantum-inspired model and calculating the similarity between them.

In graph theory -the study of graphs from a mathematical perspective-, an independent set is a set of vertices in a graph in which none of them are adjacent, that is, no two vertices have an edge that connects them. Then, given a graph *G* = (*V, E*), a maximal independent set of *G* is the largest possible independent set.

We created the graph *G* = (*V, E, L*_*V*_, *L*_*E*_), where *G* was a labelled graph representing a molecule and *V* being the set of vertices (atoms or rings in the molecule), *E* being the set of edges (bonds between atoms or rings), *L*_*V*_ being the set of labels assigned to each vertex, and *L*_*E*_ being the set of labels assigned to each edge. Labels for vertices and edges encode properties from atoms, rings, or bonds. All these features, both for vertices as well as edges, are generated using RDKit, an open-source cheminformatics software that allows working with molecules and its properties in an easy way [23].

In our case, we used the following features for vertex -atom or ring- labelling:

- Symbol.
- Number of explicit Hydrogens.
- Number of implicit Hydrogens.
- Degree of the atom (number of bonded neighbors in the graph).
- Explicit valence.
- Formal charge.
- Whether it is in a ring or not.

In particular, for ring structures we added the values from the labels of its atoms, except for the case of symbols, which are just a dictionary with symbols as keys and their repetitions as values. In the case of edges -or bonds-, we use as a feature the bond type as a float number given by RDKit, so it is easy to make calculations based on just a number instead of a string label.

After getting the features described above, we generate the graphs with NetworkX [49], a Python package for networks, in order to get an easier representation of the molecules as well as for the conflict graph we needed to generate.

### Creating the conflict graph and solving the MWIS

Our conflict graphs were generated using the following stepwise algorithms.

#### Algorithm 1. GenerateConflictGraph(*Mol1, Mol2*).

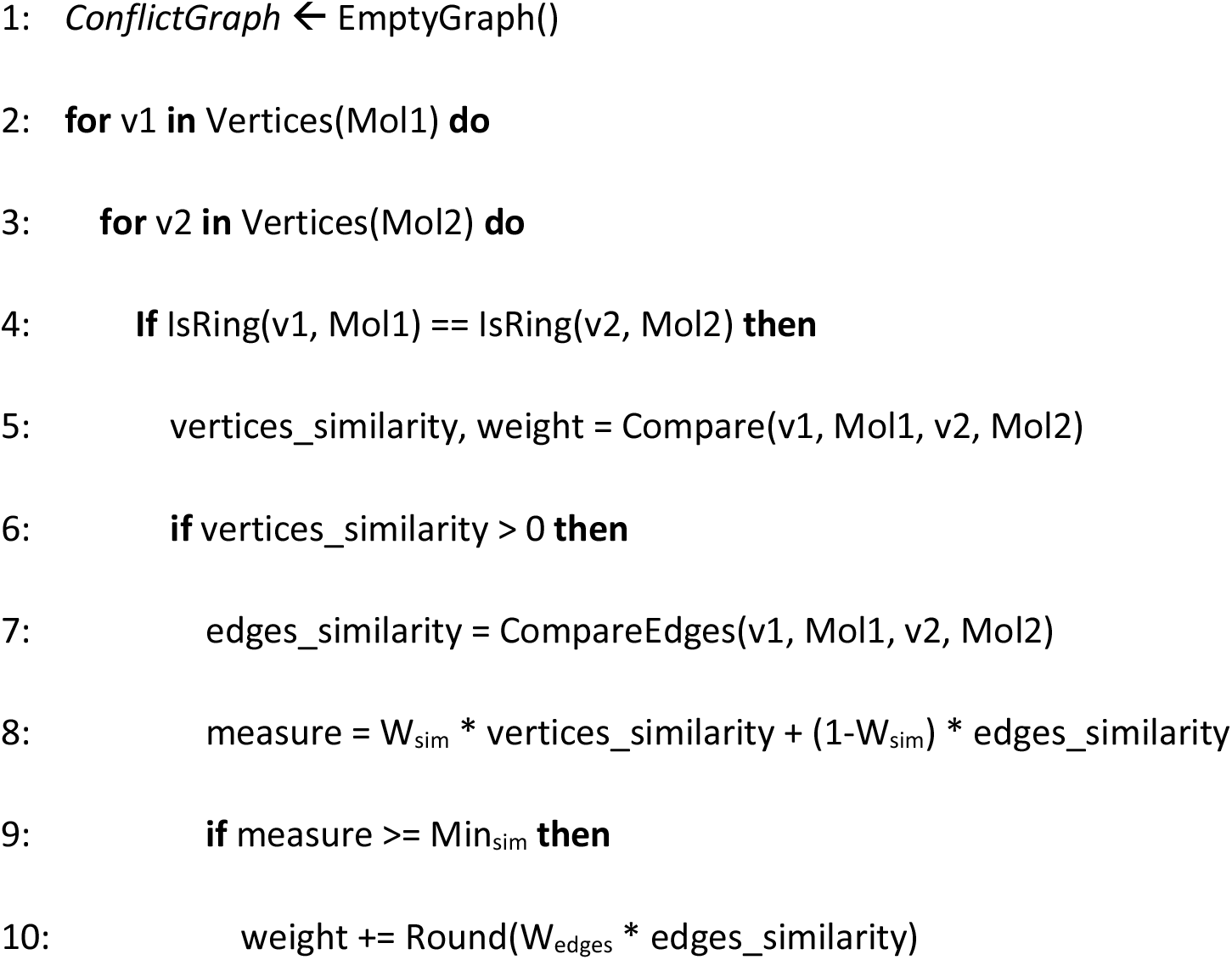
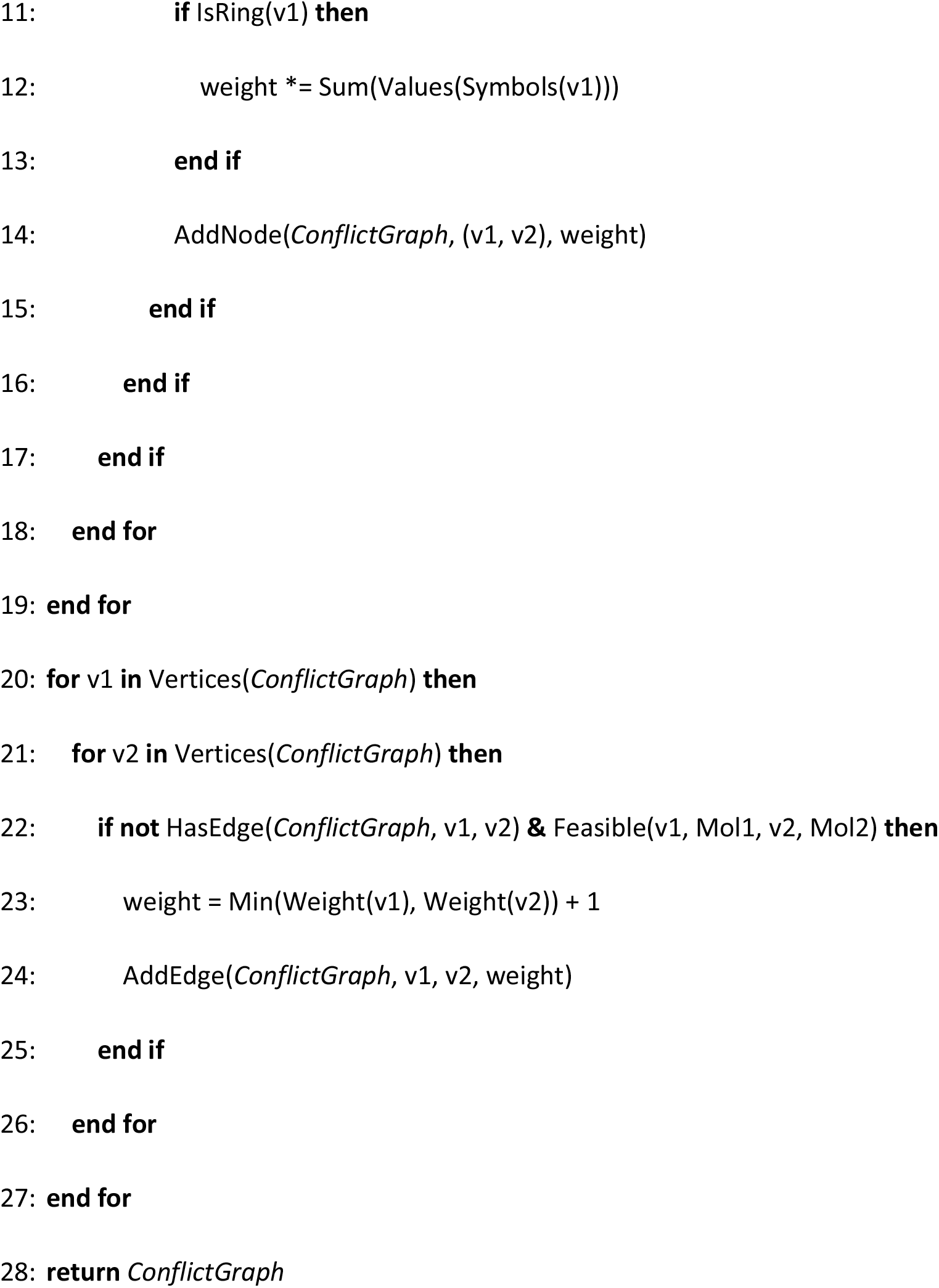

In Algorithm 1 we show the pseudo-code of the proposed method to create the conflict graph from two given molecules in the form of a graph. In particular, we traverse the set of vertices from both molecules and, if the pair of vertices are atoms or rings, we calculate its similarity in Step 5. The function *Compare* works different depending on whether the vertices are atoms or rings. Specifically, if the vertices are atoms, they must have the same symbol or belong to the halogens group (F, Cl, Br, I) in order to be compared. Then, the weight of the potential pair of vertices is the sum of the compared values of the numerical properties of atoms (number of explicit/implicit hydrogens, degree, explicit valence and formal charge). The compared value is the minimum value divided by the maximum value of each property. The similarity is the average of that value, that is, dividing the weight by the number of properties compared. On the other hand, if the vertices are rings we follow the same logic, but we need to consider all the atoms in the ring for the first filter, so two rings are compared if and only if they have exactly the same atoms.

After comparing the vertices, if they are somewhat similar (Step 6), we also compare their edges in Step 7. In particular, we compare all the adjacent vertices to the vertices being compared with the previous logic. Then, we get the average of that value. In Step 8, we get a measure of the similarity depending on the two values of similarity: the one given by the comparison of the vertices and the other one given by the similarity of their respective edges. Thus, we weigh those values differently in order to get a measure of similarity. If that measure value is higher than the minimum established value of similarity (Step 9), we add some weight to the final weight depending on the similarity of the edges (Step 10). If the vertices belong to a ring (Step 11), we also multiply this final value of weight by the number of elements in Step 12. Therefore, we consider rings heavier than atoms in our weigh method. In Step 14, we add the pair of vertices with their respective weight to the conflict graph as a new node.

When this process was finished, we needed to construct the edges among the vertices in the conflict graph. For every pair of vertices, we checked in Step 22 if they needed to be linked by an edge. We first checked they were not in the conflict graph yet and that the edge was feasible. Feasibility here means that the atoms/rings belonging to the first molecule are linked in the same way as the atoms/rings belonging to the second molecule are linked. If all the conditions were met, we calculated in Step 23 a weight for the edge. Finally, in Step 24 we added an edge with the calculated weight to the two vertices being compared, returning the newly created conflict graph in Step 28.

Once we had the conflict graph, we were ready to build the QUBO model for the optimization problem as in[38] considering that we want a minimization function instead of a maximization one:

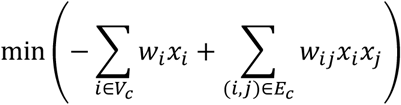

Where *x*_*i*_ is a binary variable that is equal to 1 if the vertex *i* is included in the independent set and 0 otherwise, *w*_*i*_ is the weight associated to that vertex from the conflict graph, *w*_*ij*_ is the weight associated to the vertices *i* and *j*, *V*_*c*_ is the set of vertices from the conflict graph and *E*_*c*_ is the set of edges of the conflict graph.

The first part of that expression minimizes the weights of the selected vertices from the conflict graph (the objective function) and the second part of the expression penalizes the infeasible assignments (the constraint). Building the model is trivial given the conflict graph. Since we had weights for each vertex as well as for each edge, the only thing we needed was to generate a map between vertices and binary variables for the model.

### Similarity measurement

We used the same metric as in[38] for our similarity measurement:

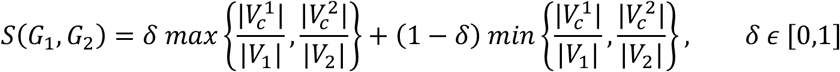

Where *G*_1_ and *G*_2_ are the original graphs from the molecules, 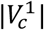 and 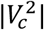 denote the number of unique vertices of *G*_1_ and *G*_2_ in the independent set of the conflict graph, │*V*_1_│ and │*V*_2_│ denote the number of vertices from *G*_1_ and *G*_2_, and *δ* is a parameter to tune the result.

Depending on the perspective, we have two different values of similarity: the similarity of *G*_1_ respect to *G*_2_ and the similarity of *G*_2_ respect to *G*_1_. Those values might be different depending on the number of similar vertices and the size of the graphs. Thus, this metric gives a value that mixes the contribution of each graph to the solution of the problem, and we were able to give more weight to one similarity value or the other one depending on the value of *δ*.

We also took into account that if the two values of similarity given by this measure (the minimum and the maximum ones) were very different, we considered the similarity as the minimum value. This high difference usually comes from two molecules very different in size, so we took the minimum value of similarity, which in this case corresponds to the bigger molecule. We set this when the maximum value is equal to or higher than the minimum one by its 50%.

### Configuration of the algorithm, similarity search and graphical representation

We implemented our algorithms in Python 3, which were run on an Intel Core i5 with 1.9 GHz and 8 GB of RAM with Microsoft Windows 10 OS for every part except for solving the mathematical model, for which we used *Digital Annealer*.

The fingerprint method was the one implemented in the RDKit library, *RDKFingerprint*, and the Tanimoto measure was calculated by the *FingerprintSimilarity* method, both of them with the default values.

The representation of QUBO similarity was done using the RDKit software.

### Cells

Vero E6 cells were kindly provided by W. Barclay (Imperial College London) and Caco-2 cells were kindly provided by C. Odendall (King’s College London). All cell lines were maintained in complete DMEM GlutaMAX (Gibco) supplemented with 10% foetal bovine serum (FBS; Gibco), 100 U/mL penicillin and 100μg/mL streptomycin and incubated at 37°C with 5% CO_2_.

### Viruses and propagation

SARS-CoV-2 Strain England 2 (England 02/2020/407073) was obtained from Public Health England. SARS-CoV-2 B.1.1.7, B.1.351 and B.1.617.2 variants isolates were kindly provided by W. Barclay (Imperial College London). Viral stocks were produced by infecting Vero E6 cells (England 02/2020/407073) or Vero E6 cells expressing TMPRSS2 (B.1.1.7, B.1.351 and B.1617.2) using an MOI of 0.01. Virus-containing supernatants were collected 72 h after infection and centrifuged at 1500 rpm for 10 min, aliquoted and stored at −80°C. The infectious virus titres were determined by plaque assay in Vero E6 cells.

### SARS-CoV-2 infection

For the drug inhibition assays, BMS-986094 (Bio-techne, UK), cobamamide (Sigma, UK), methylcobalamin (Sigma, UK), hydroxocobalamin (Caymanchem, USA) and Remdesivir (Stratech, UK) were diluted in dimethyl sulfoxide (DMSO) and added to 96-well plates of Vero E6 cells for 2 h before infection. Later, Vero E6 and Caco-2 cells were infected with SARS-CoV-2 England 02/2020/407073, B.1.1.7 or B.1.351 isolates at an MOIs of 0.05 and 0.5, respectively for 1 h. Then, the cells were washed with PBS and cultured in fresh drug-containing medium for a further 48 h. Virus production in the culture supernatants was quantified by plaque assay using Vero E6 cells and cells were collected for RNA extraction.

### RNA Extraction and real-time PCR

Total RNA was isolated from Vero E6 and Caco-2 cells 48 hours after infection using RNAdvance Viral Kit (Beckman) using a KingFisher and RNeasy Mini Kit (Qiagen), respectively. cDNA was generated using the High-Capacity cDNA Reverse Transcription Kit or H Minus RT kit (Thermo Fisher). Two regions of the viral genome of SARS-COV-2 were amplified. The first set of primers N-FW (5’-TTACAAACATTGGCCGCAAA-3’), N-RV (5’-GCGCGACATTCCGAAGAA-3’) and the probe N-probe (5’-FAM-ACAATTTGCCCCCAGCGCTTCAG-BHQ1-3’) amplified a region specific viral N RNA as a measure of total viral RNA and the second set of primers RdRP-FW: (5’-GTGARATGGTCATGTGTGGCGG-3’), RdRP-RV (5’-CARATGTTAAASACACTATTAGCATA-3’) and the probe RdRP-Probe (5’-FAM-CAGGTGGAACCTCATCAGGAGATGC-BHQ1-3’) amplified a fragment of the viral RNA-dependent RNA polymerase (RdRp) as a measure of genomic viral RNA. The fold change in viral RNA was normalized with the amplification of a fragment of human RNAse P using the primers RP-FW (5’-AGA TTTGGACCTGCGAGCG-3’), RP-RV (5’-GAG CGG CTG TCT CCA CAA GT-3’) and the probe RP-probe (5’-FAM-TTCTGACCTGAAGGCTCTGCGCG–BHQ-1-3’).

### IC_50_ calculation

The IC_50_ value was defined as the drug concentration at which there was a 50% decrease in the titre of supernatant virus. Data were analysed using Prism 9.0 (GraphPad), and IC_50_ values were calculated by nonlinear regression analysis using the dose–response (variable slope) equation.

### Citotoxicity assay

In order to assess the cytotoxicity of the compounds, Vero E6 cells or Caco-2 cells were treated 2 h before infection with the different compounds at the indicated concentrations. 48 hours after treatment, cells were incubated with 3-(4,5-dimethylthiazol-2-yl)-2,5-diphenyltetrazolium bromide (MTT) for 4 hours in the dark at 37°C with 5% CO_2_. Supernatants were then removed, and cells containing formazan were resuspended in DMSO, incubated 10 min at room-temperature and absorbance was measured to quantify cell viability.

### Statistical analysis

Results in bar charts are expressed as means ± standard deviation for experimental replicates in each case. Differences between the experimental groups were evaluated by ordinary one-way ANOVA with Dunnett’s multiple comparisons test using Prism 9.0 (GraphPad). * indicates *P* < 0.05, ** indicates *P* < 0.01, *** indicates *P* < 0.01 and **** indicates *P* < 0.0001.

## Acknowledgements

The authors thank Fujitsu Limited for providing access to *Digital Annealer* and Fujitsu Spain for all the support and commitment. This work was funded by King’s Together Rapid COVID-19 Call awards to MHM and RTMN, a Huo Family Foundation Award to MHM and RTMN, and NIAID Awards AI150472 and AI076119 to MHM. We acknowledge the Genotype-to-Phenotype UK National Virology Consortium funded by MRC/UKRI (MR/W005611/1) and Wendy Barclay’s lab at Imperial College and Public Health England for providing viral isolates. This research was funded in whole, or in part, by the Wellcome Trust (106223/Z/14/Z to MHM; 213984/Z/18/Z to RTMN). This work was supported by the Department of Health via a National Institute for Health Research comprehensive Biomedical Research Centre award to Guy’s and St. Thomas’ NHS Foundation Trust in partnership with King’s College London and King’s College Hospital NHS Foundation Trust. For the purpose of open access, the author has applied a CC BY public copyright licence to any Author Accepted Manuscript version arising from this submission.

## Author contributions

J.M. J-G., A.M. O-P acquired, analyzed and interpreted all viral work; B. M. M. designed the computational modelling, acquired and interpreted the *in silico* data; T.J.A.M. and A.R. acquired molecular data; J.I.D-H, A.M.P. and C.C.D. supervised the computational work; M.Z. provided supervision and clinical input; M.H.M. and R.T.M-N. provided supervision and designed the work. All authors approved the manuscript.

## Competing interests

The authors do not declare any competing interests related to this work.

## Notes

### Competing Interest Statement

The authors have declared no competing interest.

### Summary of Updates

New author and Acknowledgements; Author Contributions

